# Global distribution and evolution of *Mycobacterium bovis* lineages

**DOI:** 10.1101/642066

**Authors:** Cristina Kraemer Zimpel, José Salvatore L. Patané, Aureliano Coelho Proença Guedes, Robson F. de Souza, Taiana T. Silva-Pereira, Naila C. Soler Camargo, Antônio F. de Souza Filho, Cássia Y. Ikuta, José Soares Ferreira Neto, João Carlos Setubal, Marcos Bryan Heinemann, Ana Marcia Sa Guimaraes

## Abstract

*Mycobacterium bovis* is the main causative agent of zoonotic tuberculosis in humans and frequently devastates livestock and wildlife worldwide. Previous studies suggested the existence of genetic groups of *M. bovis* strains based on limited DNA markers (a.k.a. clonal complexes), and the evolution and ecology of this pathogen has been only marginally explored at the global level. We have screened over 2,600 publicly available *M. bovis* genomes and newly sequenced two wildlife *M. bovis* strains, gathering 823 genomes from 21 countries and 21 host-species, including humans, to complete a phylogenomic analyses. We propose the existence of four distinct global lineages of *M. bovis* (Lb1, Lb2, Lb3 and Lb4) underlying the current disease distribution. These lineages are not fully represented by clonal complexes and are dispersed based on geographic location rather than host species. Our data divergence analysis agreed with previous studies reporting independent archeological data of ancient *M. bovis* [(South Siberian infected skeletons at ∼2,000 years BP (before present)] and indicates that extant *M. bovis* originated during the Roman period, subsequently dispersing across the world with the discovery and settlement of the New World and Oceania, directly influenced by trades among countries.

## Introduction

Tuberculosis (TB) is the leading infectious killer in the world and approximately 10 million new cases are reported annually. In 2017, 1.6 million people died of TB and over 95% of these deaths occurred in low and middle-income countries (World Health Organization, 2018). The disease is strongly linked to poverty, with its prevalence following a socioeconomic gradient within and among countries. In addition, there is a significant, and often neglected, contributor to the global disease burden, which is the zoonotic transmission of bovine TB to humans. The WHO (World Health Organization) estimated that 142,000 new cases and 12,500 deaths occurred due to zoonotic TB in 2017 (World Health Organization, 2018), numbers that are likely underestimated due to lack of routine surveillance data from most countries (Olea-Popelka et al., 2017). People with zoonotic TB face arduous challenges; the etiologic agent is intrinsically resistant to pyrazinamide, one of the first-line drugs used in TB treatment, and a possible association with extra-pulmonary disease often delays diagnostics and treatment initiation. In addition, bovine TB results in severe economic losses for livestock producers worldwide, respecting no borders and repeatedly affecting animal conservation efforts due to the establishment of wildlife reservoirs or spillover events from cattle to associated animal populations. In order to eradicate TB by 2,030 as part of the United Nations (UN) Sustainable Development Goals, it is imperative that future prevention and control strategies focus on all forms of TB in humans, including its interface with animals.

Human and animal TB are caused by members of the *Mycobacterium tuberculosis* Complex (MTBC). The MTBC is a clonal bacterial group composed of 12 species or ecotypes with variable virulence and host tropism (Galagan, 2014). *Mycobacterium tuberculosis stricto sensu* is the main responsible for the TB numbers and is highly adapted to human hosts. On the other hand, *Mycobacterium bovis*, the causative agent of bovine TB, has a broader host range and is able to infect and cause disease in multiple host species, including humans, with variable populational persistence (Malone and Gordon, 2017). MTBC members have clonally evolved from a common ancestor with the tuberculous bacteria *Mycobacterium canettii*, and alignable regions of MTBC genomes are over 99.95% identical, with horizontal gene transfer and large recombination events considered absent. These pathogens have solely evolved through single nucleotide polymorphisms (SNPs), indels and small deletions of up to 12.7Kb, which translated into a phenotypic array of host tropism and virulence variations (Galagan, 2014; Brites et al., 2018).

Using whole-genome, SNP-based phylogenetic analyses, human-adapted MTBC have been classified into 7 lineages, with *M. tuberculosis* accounting for L1 to L4 and L7, and *Mycobacterium africanum* comprising L5 and L6. Each human-adapted MTBC lineage is associated with specific global geographical locations and lineage-associated variations in virulence, transmission capacity and in the propensity to acquire drug resistance have been reported (de Jong et al., 2010; Coscolla and Gagneux, 2014). Thus, regional prevalence of specific lineages or sub-lineages have consequences for the epidemiology of TB worldwide. A similar attempt to classify *M. bovis* into different genetic groups was made prior to the large-scale availability of whole-genome sequences and started with the identification of clonal complexes (CCs). Accordingly, four *M. bovis* CCs have been described (African 1 and 2, European 1 and 2), and these are determined based on specific deletions ranging from 806 to 14,094 bp (base pairs), SNPs and spoligotypes. As with *M. tuberculosis* lineages, *M. bovis* CCs appear to have distinct geographical distributions, with African 1 and 2 restricted to Africa, European 2 commonly found in the Iberian Peninsula, and European 1 distributed globally (Müller et al., 2009; Berg et al., 2011; Smith et al., 2011; Rodriguez-Campos et al., 2012). Although there are no studies specifically aimed at identifying differences in virulence patterns among *M. bovis* of different CCs, numerous articles report virulence variations among strains of *M. bovis* (Wedlock et al., 1999; Waters et al., 2006; Meikle et al., 2011; Wright et al., 2013; de la Fuente et al., 2015; Vargas-Romero et al., 2016), suggesting a possible link between bacterial genetic polymorphisms and disease development, as observed in *M. tuberculosis*.

Since the whole-genome sequence of the first *M. bovis* strain became available in 2003 (Garnier et al., 2003), increasing efforts have been made to sequence additional strains and use whole-genome information to tackle bovine and/or wildlife TB transmission within specific outbreaks or countries (Bruning-Fann et al., 2017; Sandoval-Azuara et al., 2017; Ghebremariam et al., 2018; Kohl et al., 2018; Lasserre et al., 2018; Orloski et al., 2018a; Price-Carter et al., 2018; Razo et al., 2018). However, no studies to date have comprehensively analyzed *M. bovis* genomes at a global scale. Few studies that have compared transboundary *M. bovis* strains analyzed bacterial isolates obtained from a reduced number of countries (n<9) and included small sample sizes (Dippenaar et al., 2017; Patané et al., 2017; Zimpel et al., 2017a; Ghebremariam et al., 2018; Lasserre et al., 2018). Nevertheless, attained results suggest that *M. bovis* strains are likely to cluster based on geographical location (Dippenaar et al., 2017; Kraemer Zimpel et al., 2017; Lasserre et al., 2018). In our previous study, we have also shown that few *M. bovis* genomes do not carry any CC genetic marker (Zimpel et al., 2017a), a phenomenon that was recently observed in *M. bovis* isolates from one cattle herd in the USA and from slaughterhouse cattle in Eritrea (Ghebremariam et al., 2018; Orloski et al., 2018b). These findings suggest that CCs are unlikely to represent the whole diversity of *M. bovis* strains, warranting further evaluation of *M. bovis* molecular lineages (Zimpel et al., 2017a; Lasserre et al., 2018).

In this study, we propose and discuss the existence of at least four distinct lineages of *M. bovis* in the world. We have screened over 2,600 publicly available *M. bovis* genomes and newly sequenced two wildlife *M. bovis* strains, gathering 823 *M. bovis* genomes from 21 countries and 21 different host-species, including humans, to complete a phylogenomic analyses. We also evaluated the evolutionary origin of *M. bovis* strains and lineages and correlated bacterial population dynamics with historical events to gain new insights into the widespread nature of bovine TB worldwide.

## Materials and Methods

### Genome sequencing of Brazilian *M. bovis* genomes

Two Brazilian *M. bovis* isolates obtained from a captive European bison (*Bison bonasus*) (Zimpel et al., 2017b), and a captive llama (*Llama glama*) (provided by the Laboratory of Bacterial Zoonosis of the College of Veterinary Medicine, University of São Paulo, Brazil) were reactivated in Stonebrink medium and a single colony was sub-cultured for DNA extraction using a previously described protocol (Zimpel et al., 2017a). DNA quality was evaluated using Nanodrop 2000c (Thermo Scientific, MA, USA) and Agilent 2100 High Sensitivity Chip DNA Bioanalyzer (Agilent Technologies, CA, USA). All procedures involving live tuberculous mycobacteria were performed in a Biosafety Level 3+ Laboratory (BSL-3+ Prof. Dr. Klaus Eberhard Stewien) located at the Department of Microbiology, Institute of Biomedical Sciences, University of São Paulo, Brazil.

Paired-end genomic libraries were constructed using TruSeq DNA PCR-free sample preparation kit (Illumina, CA, USA), and Illumina HiSeq2500 (Illumina v3 chemistry) was used to sequence the genomic library (100 bp). These procedures were performed at the Central Laboratory of High Performance Technologies in Life Sciences (LaCTAD), State University of Campinas (UNICAMP), São Paulo, Brazil. Illumina sequencing reads were deposited in the Sequence Read Archive (SRA) from the National Center for Biotechnology Information (NCBI) (accession numbers: SRR7693912 and SRR7693877).

### Selection of *M. bovis* genomes

We searched for genomes identified as “*Mycobacterium bovis”* deposited in SRA, NCBI. At the time of this selection (September 2018), the designation “*Mycobacterium tuberculosis* variant *bovis”* had not yet been applied. Accordingly, there were approximately 2,600 sequence read sets of *M. bovis* genomes deposited in this database. Genomes from *M. bovis* were initially selected if they: (i) were sequenced using Illumina technology [i.e., genomes sequenced using MinION (Oxford Nanopore Technologies, Oxford, United Kingdom) and PacBio (Pacific Biosciences, California, USA) were excluded]; (ii) presented known host and sample location; and (iii) were virulent *M. bovis* strains, i.e. not named as *M. bovis* BCG, *M. bovis* AN5, *M. bovis* Ravenel (Supplementary Figure S1). We focused on *M. bovis* genomes with complete metadata regarding location and host because our goal was to address the global distribution of the pathogen at country/continent level, as well as to provide insights into host-based associations.

From all initially selected genomes, four countries were over-represented (n>100) in the dataset and were subsequently subsampled (Table 1 and Supplementary Figure S1). Briefly, as to obtain host-species representation, first, all *M. bovis* genomes obtained from these countries that were isolated from non-bovine hosts were selected (except for cervids from the USA). From the remaining read sets from cattle and USA cervids, *M. bovis* genomes were randomly subsampled using the online sample size calculator (http://www.raosoft.com/samplesize.html), choosing an error of 5%, confidence level of 95%, and response distribution of 50% (as to maximize the sample size). A random number generator (https://www.randomizer.org) was used to randomly select genomes from these read sets. Following the principles of randomness and statistical sampling, the acquired sample should represent the whole selected population of *M. bovis* genomes available from these countries.

**Table 1.** Sampling of *Mycobacterium bovis* isolated from overrepresented countries.

| Country | Available read sets | Number of selected read sets |
| --- | --- | --- |
| Mexico | Cattle (395) and non-bovine (17) | Cattle (195) and non-bovine (17) |
| Northern Ireland | Cattle (144) and non-bovine (4) | Cattle (105) and non-bovine (4) |
| New Zealand | Cattle (192) and non-bovine (104) | Cattle (129) and non-bovine (104) |
| USA | Cattle (377), Cervid (154), others (51) | Cattle (153), Cervid (71) and others (51) |

We also selected 28 additional *M. bovis* genomes that were sequenced or identified after September 2018 (Brites et al., 2018). These isolates were from Germany (*n*=7), Ghana (*n*=5), Malawi (*n*=1), Republic of Congo (*n*=3), Russia (*n*=2), Switzerland (*n*=2), and United Kingdom (*n*=6). The same inclusion criteria described above was followed in the selection of these genomes. In the end, a total of 949 virulent *M. bovis* genomes were initially selected using the above described methods and criteria (Supplementary Table S1 and Figure S1).

### Sequencing quality criteria

FASTQ files of all 949 initially selected *M. bovis* genomes were downloaded from SRA, NCBI and trimmed using Trimmomatic (Sliding window: 5:20) (Bolger et al., 2014). To guarantee the selection of genomes with good sequence quality, the following quality criteria were considered after trimming: (i) coverage ≥15x, (ii) median read length of 70 bp, (ii) absence of low quality sequences and anomalous GC content as determined using FastQC, (iv) mapping coverage against reference genome *M. bovis* AF2122/97 (NC_002945.4) greater than 95% as indicated by using Burrows-Wheeler Aligner (Li and Durbin, 2010).

### Mapping and variant calling of reads

From the quality-approved *M. bovis* read sets, duplicated reads were removed using Picard v2.18.23 (https://github.com/broadinstitute/picard). Reads were mapped against the reference genome *M. bovis* AF2122/97 (NC_002945.4) using Burrows-Wheeler Aligner (Li and Durbin, 2010). SNPs were called using Samtools v1.9 mpileup (Li, 2011), and VarScan v2.4.3 mpileup2cns (Koboldt et al., 2012), using parameters of read depth of 7, minimum map quality and minimum base quality of 20. Generated vcf files were annotated using snpEff (Cingolani et al., 2012) based on the same reference genome, and manipulated using awk programming language to remove INDELs (insertions and deletions) and SNPs located in PE/PPE, transposases, integrases, maturase, phage and repetitive family 13E12 genes. Mixed-strain infection was defined when 15% of total SNPs was composed by heterogeneous SNPs. Read sets originating from mixed-strain cultures were excluded from downstream analyses. After sequencing quality checks, a total of 823 *M. bovis* genomes were selected for final analyses (Supplementary Figure S1).

### Phylogenetic reconstruction

A customized script in Python language (Supplementary File 1) was used to build a matrix of the polymorphic positions identified in all *M. bovis* genomes. IQ-Tree (Nguyen et al., 2015) was used to perform a maximum likelihood (ML) phylogenetic reconstruction, using 1,000 bootstrap inferences and model of TPM3u+F+ASC+R9. Additionally, a phylogenetic tree was constructed based on maximum parsimony (MP) with standard bootstrap (1,000 replicates) using MPBoot (Hoang et al., 2018). Both phylogenies were rooted using *M. tuberculosis* H37Rv and topology was annotated using FigTree v1.4.3 (http://tree.bio.ed.ac.uk/software/figtree/) and Mind the Graph (https://mindthegraph.com). In order to compare if the ML and MP trees were significantly similar, the Robinson-Foulds distance (Robinson and Foulds, 1981) between them was obtained, then tested for significance of this estimated distance by generating 1,000 random trees in IQ-Tree with the same number of taxa used in the analysis. Finally, the original distance to the distribution of distances among all random trees (for a total of 499,500 comparisons) was compared, and results were plotted in a histogram done in R software 3.5.0.

### SNP markers

Unique SNPs were manually searched using the generated SNP matrix described above. The SNP positions in the genomes were retrieved using a customized Python script (Supplementary File 1) and were annotated based on *M. bovis* AF2122/97 (NC_002945.4).

### Spoligotyping and Clonal Complexes

Spoligotypes of all selected genomes were investigated using SpoTyping (Xia et al., 2016). Identified genetic spacers were processed in the *Mycobacterium bovis* Spoligotype Database (www.mbovis.org) to retrieve spoligotype pattern and SB number. Intact regions of RdEu1 (European 1 - Eu1, RD17, 1,207 bp), SNP in the *guaA* gene (European 2 - Eu2), RDAf1 (African 1 - Af1, 350 bp) and RDAf2 (African 2 - Af2, 451 bp) (Müller et al., 2009; Berg et al., 2011; Smith et al., 2011; Rodriguez-Campos et al., 2012) were searched against all genomes using the BLASTn tool available in SRA, NCBI. Genomes with “no significant similarity found” as result in BLASTn when using intact regions of CC markers were mapped against the *Mycobacterium tuberculosis* H37Rv (NC_000962.3), using CLC Genomics Workbench 11 (QIAGEN, Venlo, Netherlands), to confirm the presence of RDs/SNPs.

### Population structure

Population structure was evaluated using Principal Component Analysis (PCA) based on the generated SNP matrix and using function *princomp* in R software 3.5.0 (Mardia et al., 1979; Venables and Ripley, 2002).

### Dating divergences

Molecular dating of divergences was performed using BEAST v1.10. (Suchard et al., 2018) under a coalescent approach (extended Bayesian skyline model). The GTR+G model was implemented, altogether with ascertainment bias correction (important since only SNPs are included in the alignment). The ML tree was fixed throughout the runs. A model of uncorrelated rates across branches was implemented. Molecular rates of evolution were set within the range of a Uniform [1 x 1e-6; 2 x 1e-4] s/SNP-s/b/y (substitutions/SNP_site/branch/year), based on previous studies (Ford et al., 2011; Pepperell et al., 2013; Eldholm et al., 2015; Kay et al., 2015; Lillebaek et al., 2016). The minimum age was set to 800 BP, after reasonable carbon-dating skeleton data, and to a soft upper maximum of ∼7,000 BP (a conservative upper bound), both based on Bos et al. (Bos et al., 2014) posterior estimates, under a negative exponential prior. Such a soft upper bound allows older dates in our analyses, if these are likely to occur in the posterior distribution of our estimates, therefore minimizing possible biases related to an overly conservative time range. A total of two runs were executed, each with 1 billion generations. Convergence and effective sample sizes (ESSs) were monitored in Tracer v1.7 (http://tree.bio.ed.ac.uk/software/tracer), logcombiner (within the BEAST package) was used to join results from the two runs, and treeannotator (also within the same package) was used to construct the time tree after disregarding the burnin region from each run.

## Results and Discussion

### *Mycobacterium bovis* genomes and host adaptation

After screening ∼2,600 publicly available *M. bovis* genomes using pre-determined inclusion criteria, a total of 823 virulent *M. bovis* genomes from 21 countries and 21 different host species were selected for this study (Table 2). This sample constitutes the most comprehensive global *M. bovis* dataset ever analyzed, and includes 724 genomes published previously, and two newly sequenced genomes from this study (Supplementary Table S2). We used 17,302 nucleotide polymorphic positions detected in these genomes to construct phylogenetic trees inferred with maximum likelihood (ML) and maximum parsimony (MP) methodologies (Figure 1; Supplementary Files 2 and 3 and Figures S2 and S3). Both phylogenetic methods employed (ML and MP) generated highly similar trees (Supplementary Figures S2 and S3). This observation is supported by the Robinson-Foulds distance between both trees (= 678) compared to all other comparisons among 1,000 random trees with the same number of taxa (for a total of 499,500 distances obtained), for which the minimum was 1,632, and the maximum was 1,642 (Supplementary Figure S4). Therefore, results are shown for ML only (Figures 1-4).

**Table 2.** *Mycobacterium bovis* genomes selected for analysis.

| Country | Number of genomes | Host species (# of selected read sets) |
| --- | --- | --- |
| Brazil | 3 | Cattle (1), Bison (1), Llama (1) |
| Canada | 6 | Cattle (5), Elk (1) |
| China | 2 | Chimpanzee (2) |
| Eritrea | 12 | Cattle (12) |
| Ethiopia | 2 | Cattle (2) |
| France | 2 | Cattle (1), Wild boar (1) |
| Germany | 6 | Human (6) |
| Ghana | 3 | Human (3) |
| Great Britain | 12 | Human (11), Cattle (1) |
| Italy | 2 | Human (2) |
| Mexico | 192 | Cattle (175), Human (17) |
| New Zealand | 232 | Cattle (129), Ferret (38), Pig (7), Cervid (4),<br>Stoat (1), Possum (53) |
| Northern Ireland | 88 | Cattle (84), Badger (4) |
| Panama | 2 | Cattle (2) |
| Russia | 1 | Human (1) |
| South Africa | 11 | Cattle (1), Lion (2), Kudu (1), Buffalo (7) |
| Spain | 7 | Cattle (7) |
| Switzerland | 2 | Unknown host* |
| Tunisia | 1 | Human (1) |
| United States | 233 | Cattle (124), Coyote (1), Human (2), Cervid (64),<br>Cat (7), Raccoon (12), Wild boar (6), Opossum<br>(9), Jaguar (1), Bobcat (1), Bison (1), Elk (5) |
| Uruguay | 4 | Cattle (4) |
| **Total** | **823** |  |

\**Mycobacterium bovis* genomes sequenced by Brites et al. (Brites et al., 2018).

**Figure 1.**
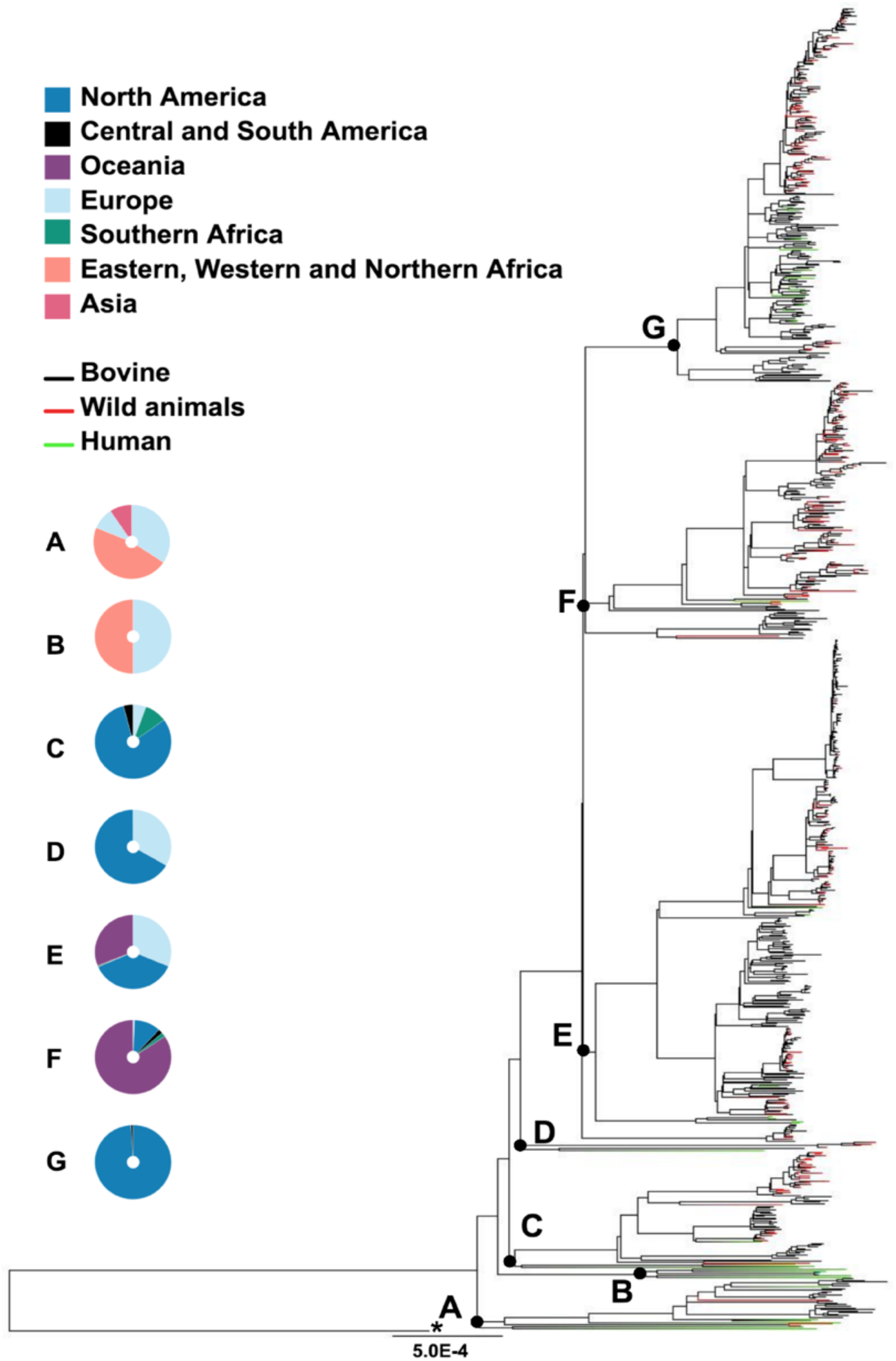
Host species and geographical distribution of *Mycobacterium bovis* isolates. Maximum likelihood phylogenetic tree based on single nucleotide polymorphisms (SNPs) of 823 *Mycobacterium bovis* genomes, using \**Mycobacterium tuberculosis* H37Rv as outgroup. The host species are marked in black tips for bovine, red tips for wild animals, green tips for humans, and in blue tips are two genomes with unknown host identification. Geographical locations of the isolates emerging from main nodes A-G (black dots) are colored in donut charts (which do not represent ancestral area reconstruction at nodes): Blue - North America; Black - South and Central America; Purple - Oceania; Pink - Europe; Green - Southern Africa; Orange - Eastern, Western and Northern Africa; Light blue - Asia. Phylogenetic tree was generated using IQ-Tree with 1,000 bootstrap values and annotated using FigTree v1.4.3 and Mind the Graph. Bootstrap values of discussed nodes are all ≥ 95% and can be visualized in Supplementary Figure S2 and Supplementary File 2. Bar shows substitutions per nucleotide.

**Figure 2.**
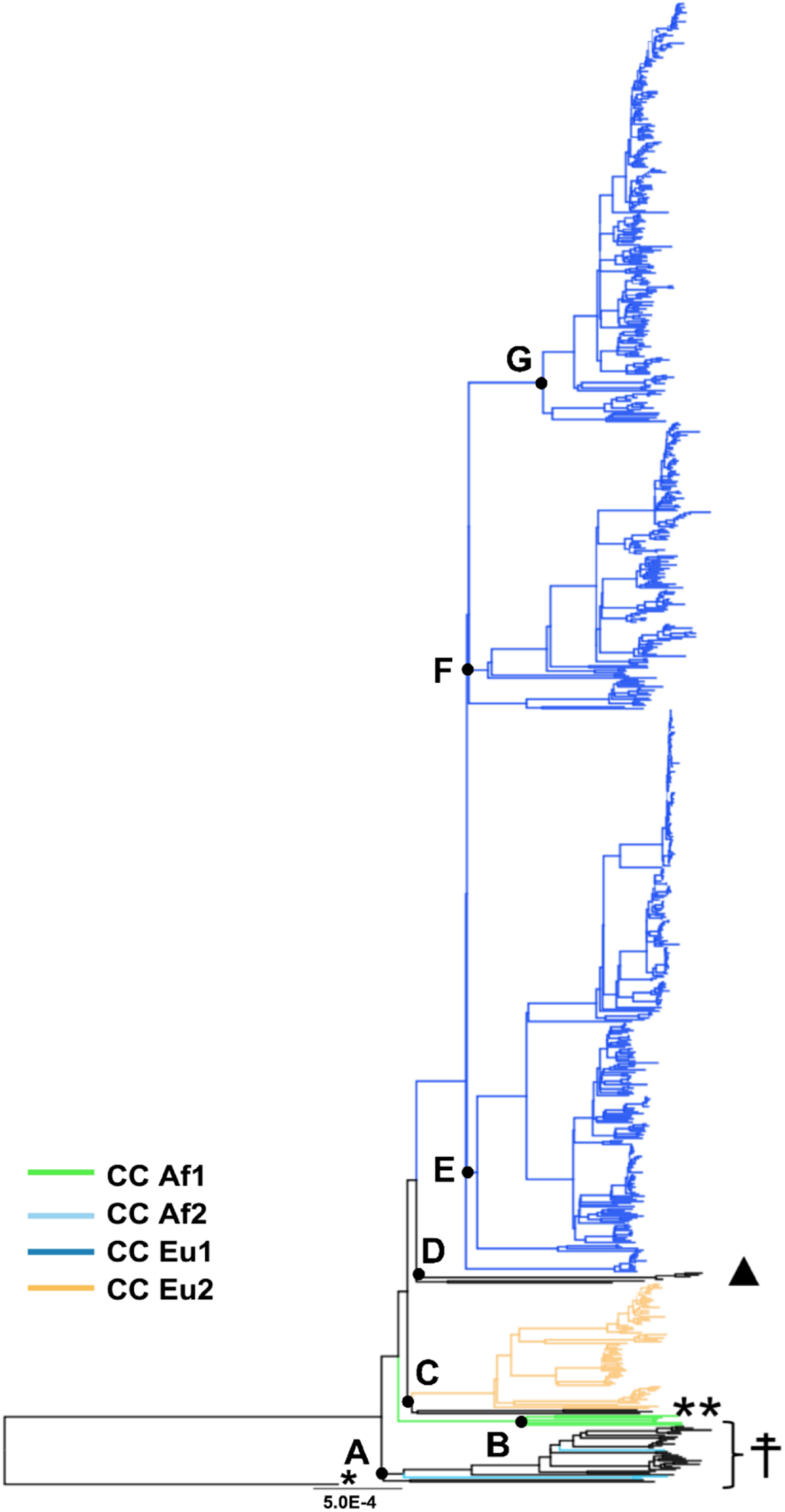
Clonal complexes characterization. Maximum likelihood phylogenetic trees based on single nucleotide polymorphisms (SNPs) of 823 *Mycobacterium bovis* genomes, using \**Mycobacterium tuberculosis* H37Rv as outgroup. Phylogenetic tree colored based on clades and clonal complexes (CC). Clonal complexes (tips): African 1 in green, African 2 in light blue, European1 in dark blue, and European 2 in yellow. Genomes without clonal complexes markers are shown in black. ☨: Cluster containing three *M. bovis* genomes of CC African 2 and 29 *M. bovis* genomes without CC markers. ** Cluster containing three *M. bovis* genomes without CC markers (black tips), but closely related to *M. bovis* of Eu2 CC marked in yellow. ▲: Cluster containing six *M. bovis* genomes without CC markers (black tips), but closely related to *M. bovis* of CC Eu1 marked in dark blue. Black dots: main nodes A-G. Phylogenetic tree was generated using IQ-Tree with 1,000 bootstrap values and annotated using FigTree v1.4.3 and Mind the Graph. Bootstrap values of discussed nodes are all ≥ 95% and can be visualized in Supplementary Figure S2 and Supplementary File 2. Bar shows substitutions per nucleotide.

**Figure 3.**
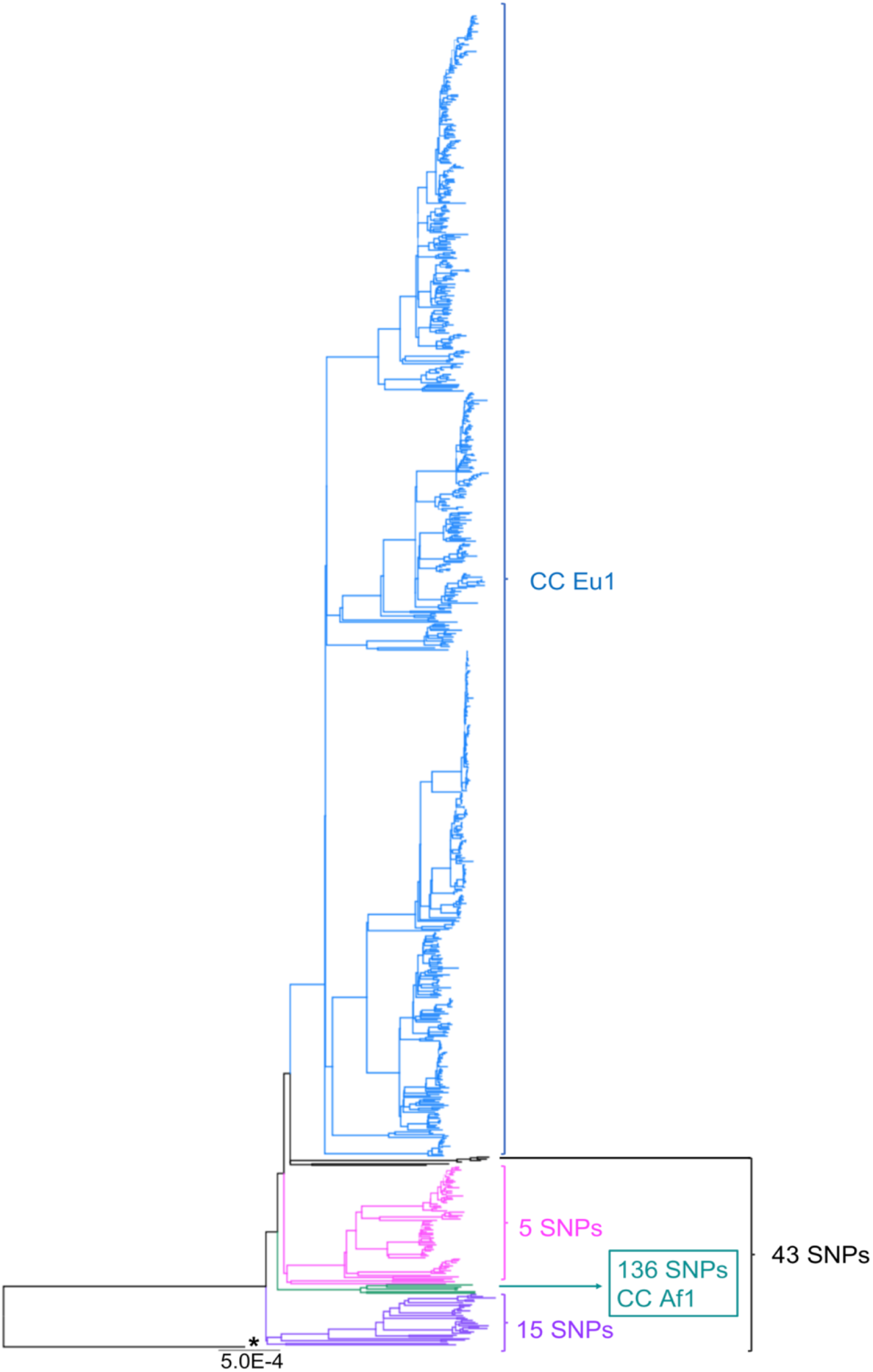
SNP (single nucleotide polymorphism) markers of *Mycobacterium bovis*. Maximum likelihood phylogenetic trees based on single nucleotide polymorphisms (SNPs) of 823 *Mycobacterium bovis* genomes, using \**Mycobacterium tuberculosis* H37Rv as outgroup. Clonal complexes: African 1 (Af1) and European 1 (Eu1). Phylogenetic tree was generated using IQ-Tree with 1,000 bootstrap values and annotated using FigTree v1.4.3. Bootstrap of discussed nodes are all ≥ 95% and can be visualized in Supplementary Figure S2 and Supplementary File 2. Bar shows substitutions per nucleotide.

**Figure 4.**
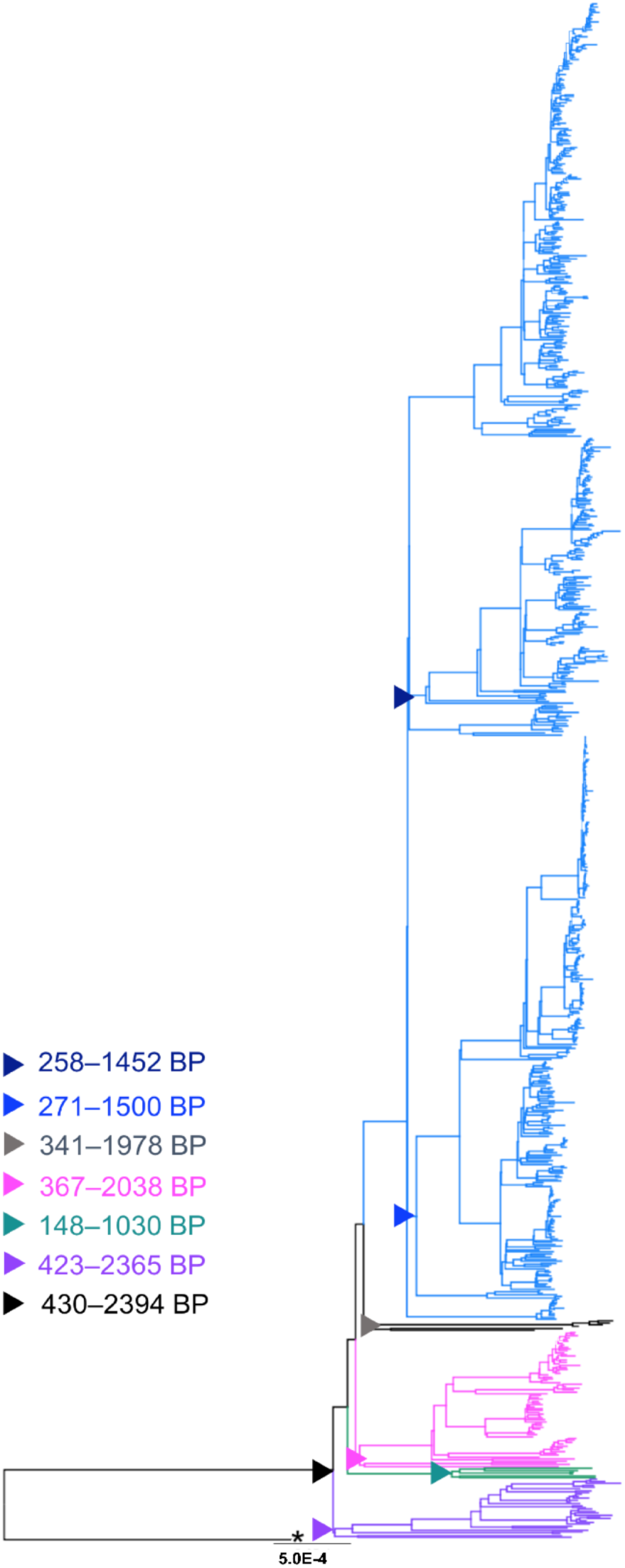
*Mycobacterium bovis* lineages and evolutionary data. Maximum likelihood phylogenetic tree of 823 *M. bovis* genomes colored based on the proposed Lineages: Lb1 - purple; Lb2 - green; Lb3 - pink; Lb4 - blue; uncharacterized genomes - grey. Estimated divergence results in years BP (Before Present) of the main nodes are marked with triangles. Outgroup: \**Mycobacterium tuberculosis* H37Rv. Phylogenetic tree was generated using IQ-Tree with 1,000 bootstrap values and annotated using FigTree v1.4.3. Bootstrap of discussed nodes are all ≥ 95% and can be visualized in Supplementary Figure S2 and Supplementary File 2. Bar shows substitutions per nucleotide.

In the generated phylogenetic tree, host species are found dispersed among different clades (Figure 1 - colored tips), while most *M. bovis* genomes cluster according to specific geographic locations (Figure 1). This finding suggests that geographic proximity between hosts and their contact rates has played a more important role in determining host range of *M. bovis* than phylogenetic distance among hosts. The lack of host clustering supports the hypothesis of *M. bovis* being a generalist member of MTBC, able to infect different host species irrespective of the bacterial genetic makeup. Interestingly, all *M. bovis* genomes originating from the African Continent, except for South Africa, appeared as originating from relatively older nodes of the phylogenetic tree (Figure 1 - isolates emerging from nodes A and B), suggesting the continent is a possible center of dispersion.

### Clonal complexes do not represent the whole diversity of *M. bovis* genomes

We have searched for CC genetic markers (Eu1 - European 1, Eu2 - European 2, Af1 - African 1, Af2 - African 2) in all selected genomes using sequence similarity detection approaches. The distribution of CCs onto the phylogenetic tree is shown in Figure 2. Accordingly, 38 out of the 823 (4.6%) analyzed *M. bovis* genomes do not have genetic markers of any of the four CCs. The remaining *M. bovis* isolates were found to cluster according to their CC classification. The vast majority (29/38, 76.32%) *of M. bovis* genomes not classified within CCs appeared dispersed, emerging from relatively ancestral nodes (Figure 2). This finding suggests that extant isolates clustering in ancient nodes were probably not included in prior studies evaluating CCs and have thus been poorly studied until now.

To facilitate data presentation and interpretation, we describe below the CC classification and country of origin of the analyzed *M. bovis* isolates in order from the most ancient to the most recent nodes of the phylogenetic tree depicted in Figure 2. Accordingly, three *M. bovis* isolates of the CC Af2 (Figure 2 - light blue) emerged from the most basal node (node A) of the phylogenetic tree together with 29 *M. bovis* genomes without CC classification (Figure 2 - indicated as ☨, isolates emerging from node A). These three *M. bovis* genomes of CC Af2 comprise one isolate from a wild boar in France and two isolates from chimpanzees in China. The 29 closely-related *M. bovis* genomes without clonal complex classification were obtained in Africa [Eritrea (*n*=12), Ethiopia (*n*=2), Tunisia (*n*=1)], Europe [Italy (*n*=2), Spain (*n*=7), Germany (*n*=1)], Russia (*n*=1) and the USA (*n*=3). In a subsequent cluster (Figure 2 - isolates emerging from node B), it is possible to observe six *M. bovis* isolates classified as CC Af1 obtained in Africa [Ghana (*n*=3)] and Europe [Germany (*n*=1), Switzerland (*n*=2)] (Figure 2 - in green).

To date, *M. bovis* strains of the CCs Af1 and Af2 have only been described in Africa (Berg et al., 2011; Firdessa et al., 2013). The finding of *M. bovis* strains of CC Af2 infecting chimpanzees in China may be explained by pathogen importation from the African continent into that country or, less likely, spill-over from infected livestock to captive animals. A thorough characterization of *M. bovis* strains from China in the future could elucidate which CC is more frequent in that country. On the other hand, the presence of both African CCs in countries of Europe, infecting a wild boar (France), humans (Germany) and unknown hosts (Switzerland), is interesting and warrants further investigation into the actual origin of these isolates. Ten out of the 43 *M. bovis* genomes (23.26%) described in the paragraph above originated from isolates obtained from humans, being four from Africa (Ghana=3; Tunisia=1), four from Europe (Germany=2; Italy=2), one from Russia, and one from the USA. Unfortunately, demographic characteristics of these patients are not described in the related literature (Casali et al., 2014; Walker et al., 2015; Brites et al., 2018; Kandler et al., 2018). With current bovine TB control, the zoonotic transmission of *M. bovis* in Europe and the USA is considered rare. Thus, we speculate that European and North American cases in humans may have arisen from zoonotic transmission acquired in the past that appears in elderly people (Majoor et al., 2011; Davidson et al., 2017) and/or imported human cases of zoonotic TB from countries where bovine TB is highly endemic.

The phylogenetic tree in Figure 2 also shows 70 *M. bovis* genomes carrying the SNP marker of the CC Eu2 (Figure 2 - in yellow, part of isolates emerging from node C). These isolates were obtained from the Americas [Brazil (*n*=3), Canada (*n*=2), Mexico (*n*=17), USA (*n*=40)], Africa [South Africa (*n*=7)], and Europe [Germany (*n*=1)]. There were only three *M. bovis* genomes without CC classification isolated from humans in Germany that appeared closely related to genomes of the CC Eu2 (Figure 2 - indicated as **, part of isolates also emerging from node C). The CC Eu2 has been described as dominant in the Iberian Peninsula and also detected in other countries of Europe and in Brazil (Rodriguez-Campos et al., 2012; Zimpel et al., 2017a). Its actual worldwide occurrence, however, is unknown.

The majority of the genomes in the dataset (706/823, 85.8%) were classified as Eu1 (i.e. having the deletion RDEu1) and originated from the Americas [USA (*n*=186), Mexico (*n*=175), Uruguay (*n*=4), Canada (*n*=4), Panama (*n*=2)], Oceania [New Zealand (*n*=88)], Europe [Great Britain (*n*=11), Northern Ireland (*n*=232)] and Africa [South Africa (*n*=4)] (Figure 2 - dark blue, isolates emerging from nodes E, F and G). Interestingly, *M. bovis* of the CC Eu1 have been found to be dispersed worldwide (Smith et al., 2011), and herein, they emerged from the most recent evolutionary node of the phylogenetic tree. There were also six genomes closely and basally related to *M. bovis* genomes of CC Eu1 that did not carry any CC marker and were isolated in Europe [France (*n*=1), Great Britain (*n*=1)] and the USA (*n*=4) (Figure 2 - indicated as ▲, isolates emerging from node D).

### SNP markers

We used the generated SNP matrix to search for SNPs that were unique to specific groups of genomes of the phylogenetic tree (i.e. present in 100% of the strains in the analyzed group and not present in strains outside of that group) (Figure 3 and Supplementary Table S3). A total of 15 SNPs were found to be unique to the group comprising the three genomes of *M. bovis* classified as CC Af2 and the 29 closely-related *M. bovis* genomes without CC classification (i.e. the group originating from the most basal node of the phylogenetic tree) (Figure 3 - in purple). Therefore, these 15 SNPs are more stable markers of this group than the RDAf2 deletion. On the other hand, 136 SNPs were found to be unique to the small group of six *M. bovis* genomes with the RDAf1 deletion, further supporting the segregation of this phylogenetic group (Figure 3 - in green).

In addition to the SNP in *guaA* gene that determines the CC Eu2, 81 SNPs were found to be unique to *M. bovis* genomes classified as CC Eu2 in this study (data not shown). We also found 5 SNPs that were unique to the *M. bovis* of CC Eu2 and the three closely-related *M. bovis* genomes without CC classification, supporting the genetic segregation of this phylogenetic group (Figure 3 - in pink).

There were no SNP markers unique to *M. bovis* of the CC Eu1; the only stable genetic marker is the deletion RDEu1. The six genomes without CC classification that are closely related to *M. bovis* genomes of CC Eu1 did not share unique SNPs in common and with genomes of the CC Eu1. Interestingly, these genomes shared 43 unique SNPs with the most basal genetic groups of the phylogenetic tree. SNPs’ genomic positions and annotations are reported in Supplementary Table S3.

### A proposal for global lineages of *M. bovis*

Based on the observed tree topology, CC distribution, geography, and SNP markers, we propose the existence of at least four major global lineages of *M. bovis*, which we define as Lb1, Lb2, Lb3 and Lb4 (Figure 4 and Table 3). Lineage Lb1, emerging from the most basal node, is composed of 32 *M. bovis* isolates, encompassing the three representatives of CC Af2 and the 29 closely-related genomes without CC classification (Figure 4 - in purple). We provisionally propose the 15 above-described unique SNPs as identification markers for this lineage (Supplementary Table S3). The observed geographical origin of extant Lb1 isolates points towards North and East Africa and Europe, although it can also be detected, at a lower frequency, in the United States (three Lb1 isolates out of 233 *M. bovis* isolates from the USA). Lb2 lineage (Figure 4 - in green) is composed of the six *M. bovis* genomes of CC Af1. These genomes are robustly segregated by the RDAf1 deletion and 136 SNP markers. It is important to highlight, however, that further genomes should be sequenced to fully characterize this lineage. The extant Lb2 comprise isolates from Ghana (n=3), Germany (n=1) and Switzerland (n=2). Unfortunately, host information was not available for the isolates obtained in Switzerland. It is possible that this *M. bovis* strain was isolated from a human patient, which precludes geographical origin analyses due to human migration. Nevertheless, both extant Lb1 and Lb2 have strong ties to North, East and West Africa and to a lesser degree with Europe.

**Table 3.** Distribution of *Mycobacterium bovis* lineages in the studied countries.

| Country | Lb1 | Lb2 | Lb3 | Lb4 |
| --- | --- | --- | --- | --- |
| Eritrea | 12 | - | - | - |
| Ethiopia | 2 | - | - | - |
| Tunisia | 1 | - | - | - |
| France | 1 | - | - | - |
| Germany | 1 | 1 | 4 | - |
| Spain | 7 | - | - | - |
| Italy | 2 | - | - | - |
| China | 2 | - | - | - |
| Russia | 1 | - | - | - |
| Ghana | - | 3 | - | - |
| Switzerland | - | 2 | - | - |
| United States | 3 | - | 40 | 186 |
| Mexico | - | - | 17 | 175 |
| Canada | - | - | 2 | 4 |
| Brazil | - | - | 3 | - |
| South Africa | - | - | 7 | 4 |
| Northern Ireland | - | - | - | 88 |
| Great Britain | - | - | - | 11 |
| New Zealand | - | - | - | 232 |
| Panama | - | - | - | 2 |
| Uruguay | - | - | - | 4 |
| **Total \*** | **32** | **6** | **73** | **706** |

Lb - Lineage of *Mycobacterium bovis*. \*From 823 studied genomes, six were not included in the referred lineages.

Lb3 is composed of the 70 *M. bovis* of CC Eu2 and the three genomes without CC classification (Figure 4 - in pink), being supported by five unique SNPs (Supplementary Table S3). Interestingly, *M. bovis* genomes of CC Eu2 comprises a rapidly evolving and geographically diverse sublineage (Lb3.1) when compared to the three genomes obtained from humans in Germany (Lb3.2). Again, geographic origin and country history of these infected humans are not described in the related literature (Walker et al., 2015) and it is possible that these patients acquired the infection in the past, as described above, representing isolates of an older origin and/or in another country. The CC Eu2 is not a stable marker for Lb3 as a whole; instead, the five unique SNPs may be used as identification markers for Lb3 (Supplementary Table S3).

Finally, lineage Lb4 (Figure 4 - in blue) is composed of 706 genomes of *M. bovis* of the CC Eu1, indicating that RDEu1 is a stable marker of this evolutionarily recent lineage. Lineage Lb4 is composed of a higher number of genomes when compared to other lineages due to over-representation of geographically-restricted genomes from United States, New Zealand, Northern Ireland and Mexico. The ladder-like tips observed in Lb4 are consistent with recent population expansion across subpopulations of Lb4, which may be directly influenced by the persistence of *M. bovis* in different wildlife populations, variations in the efficiency of bovine TB control programs and geographic isolation. The tree topography suggests that once an ancestral *M. bovis* strain was introduced into each of these countries, this subpopulation clonally expanded in geographic isolation. This finding agrees with cattle trade and export patterns from Europe and Africa into the New World, Oceania, and Northern Ireland, which eventually led to the introduction of bovine TB in these regions.

With the described SNPs and deletions, it is not possible to determine whether the remaining six closely-related genomes without a CC marker (from France, Great Britain, and the USA) can be classified within a specific lineage (Figure 4 - in black). These genomes do not share unique genetic markers with Lb4 or Lb3 (i.e. SNPs not present in any other lineage), but do share 43 SNPs with the more ancient lineages Lb1, Lb2 and Lb3 (Figure 3). Further genomes should be sequenced to better characterize this group.

### Spoligotyping patterns are correlated with *M. bovis* lineages

To further support our findings related to the global *M. bovis* lineages, we also evaluated the spoligotype of all isolates. There was a good association between spoligotype and the four *M. bovis* lineages (Table 4), except for SB0265 and SB1345, which appear in both Lb3 and Lb4. Thus, the vast majority of the patterns were specific to the predicted lineages, demonstrating that spoligotyping can be a supportive tool to infer these groups, albeit keeping in mind that homoplasy is a common phenomenon in spoligotypes (i.e. identical spoligotype patterns can occur independently in unrelated lineages because the loss of spacer sequences is a common event) (Warren et al., 2002) and may also occur with lineage classification (e.g. SB0265 and SB1345 appear in Lb3 and Lb4).

**Table 4.**
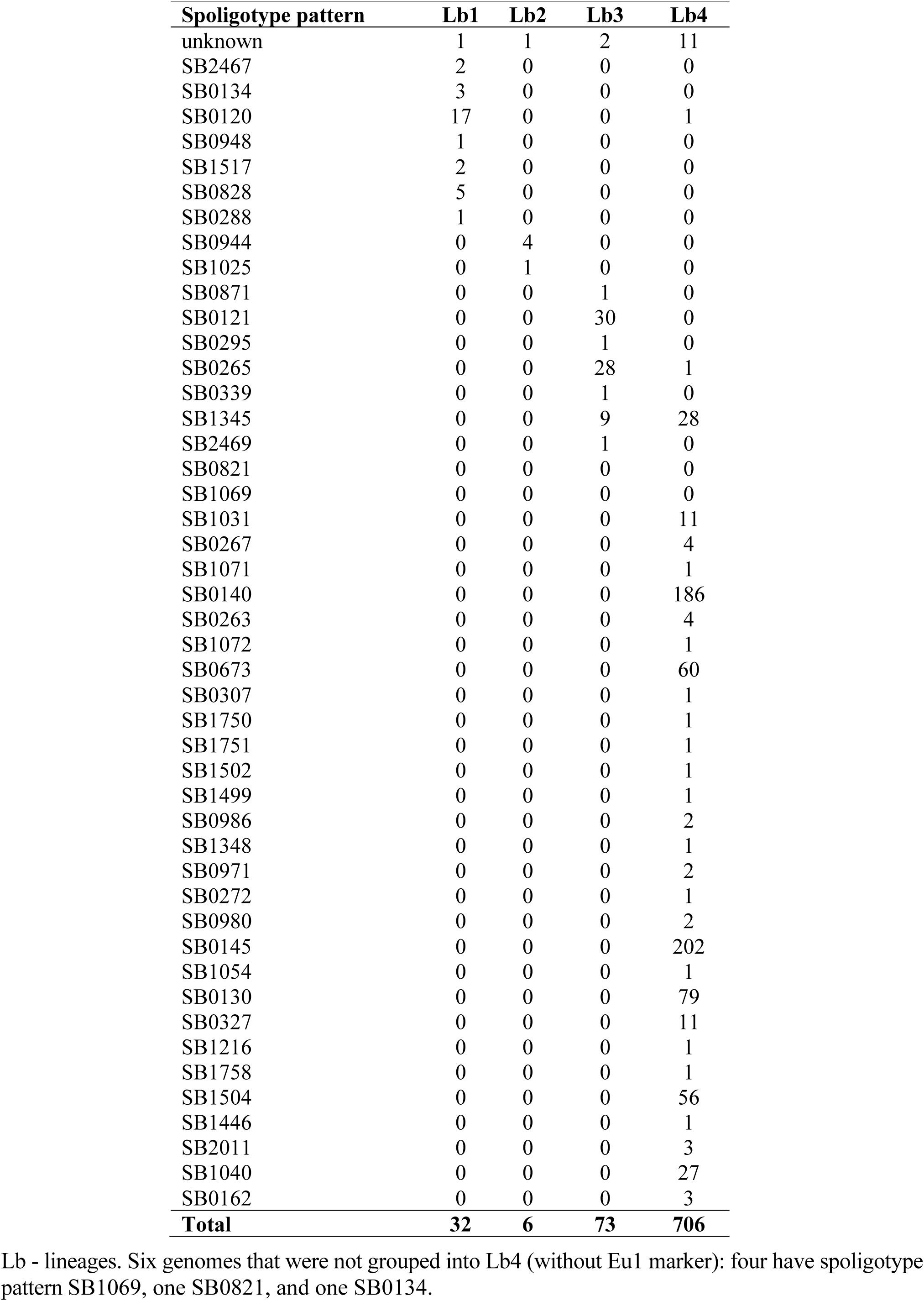
Spoligotype patterns of *Mycobacterium bovis* strains separated by lineages.

| **Spoligotype pattern** | **Lb1** | **Lb2** | **Lb3** | **Lb4** |
| --- | --- | --- | --- | --- |
| unknown | 1 | 1 | 2 | 11 |
| SB2467 | 2 | 0 | 0 | 0 |
| SB0134 | 3 | 0 | 0 | 0 |
| SB0120 | 17 | 0 | 0 | 1 |
| SB0948 | 1 | 0 | 0 | 0 |
| SB1517 | 2 | 0 | 0 | 0 |
| SB0828 | 5 | 0 | 0 | 0 |
| SB0288 | 1 | 0 | 0 | 0 |
| SB0944 | 0 | 4 | 0 | 0 |
| SB1025 | 0 | 1 | 0 | 0 |
| SB0871 | 0 | 0 | 1 | 0 |
| SB0121 | 0 | 0 | 30 | 0 |
| SB0295 | 0 | 0 | 1 | 0 |
| SB0265 | 0 | 0 | 28 | 1 |
| SB0339 | 0 | 0 | 1 | 0 |
| SB1345 | 0 | 0 | 9 | 28 |
| SB2469 | 0 | 0 | 1 | 0 |
| SB0821 | 0 | 0 | 0 | 0 |
| SB1069 | 0 | 0 | 0 | 0 |
| SB1031 | 0 | 0 | 0 | 11 |
| SB0267 | 0 | 0 | 0 | 4 |
| SB1071 | 0 | 0 | 0 | 1 |
| SB0140 | 0 | 0 | 0 | 186 |
| SB0263 | 0 | 0 | 0 | 4 |
| SB1072 | 0 | 0 | 0 | 1 |
| SB0673 | 0 | 0 | 0 | 60 |
| SB0307 | 0 | 0 | 0 | 1 |
| SB1750 | 0 | 0 | 0 | 1 |
| SB1751 | 0 | 0 | 0 | 1 |
| SB1502 | 0 | 0 | 0 | 1 |
| SB1499 | 0 | 0 | 0 | 1 |
| SB0986 | 0 | 0 | 0 | 2 |
| SB1348 | 0 | 0 | 0 | 1 |
| SB0971 | 0 | 0 | 0 | 2 |
| SB0272 | 0 | 0 | 0 | 1 |
| SB0980 | 0 | 0 | 0 | 2 |
| SB0145 | 0 | 0 | 0 | 202 |
| SB1054 | 0 | 0 | 0 | 1 |
| SB0130 | 0 | 0 | 0 | 79 |
| SB0327 | 0 | 0 | 0 | 11 |
| SB1216 | 0 | 0 | 0 | 1 |
| SB1758 | 0 | 0 | 0 | 1 |
| SB1504 | 0 | 0 | 0 | 56 |
| SB1446 | 0 | 0 | 0 | 1 |
| SB2011 | 0 | 0 | 0 | 3 |
| SB1040 | 0 | 0 | 0 | 27 |
| SB0162 | 0 | 0 | 0 | 3 |
| **Total** | **32** | **6** | **73** | **706** |

Lb - lineages. Six genomes that were not grouped into Lb4 (without Eu1 marker): four have spoligotype pattern SB1069, one SB0821, and one SB0134.

### Principal component analysis

To further evaluate the genetic relationship among *M. bovis* lineages, the SNP matrix was subjected to a principal component analysis (Figure 5). Results suggest a robust segregation of the modern lineage Lb4 and a close genetic relatedness of the more ancient lineages Lb1, Lb2 and Lb3. These findings directly reflect the results of the phylogenetic tree and SNP markers, as the most basal clusters appeared more closely related and shared 43 unique SNPs. The resulting PCA analysis is similar to what is observed using an equivalent approach with *M. tuberculosis* lineages, in which the modern lineages of *M. tuberculosis* (L2, L3 e L4) appear closely related, whereas the ancient lineages (L1, L5, L6 and L7) are found segregated from the rest, in different groups (Brites et al., 2018). It also suggests that Lb4, which is markedly characterized by the Eu1 deletion, is likely composed of three sublineages that correspond to *M. bovis* genomes emerging from nodes E, F, G of the phylogenetic tree depicted in Figure 1. The *M. bovis* genomes emerging from node G are mostly from North America (two genomes only are from Panama and Uruguay), which may indicate that these Lb4 sublineages have been evolving under geographical segregation. Our findings highlight the need to further evaluate the virulence phenotype of the proposed *M. bovis* lineages, as ancient and modern *M. tuberculosis* lineages display distinct abilities to cause disease (de Jong et al., 2010; Portevin et al., 2011).

**Figure 5.**
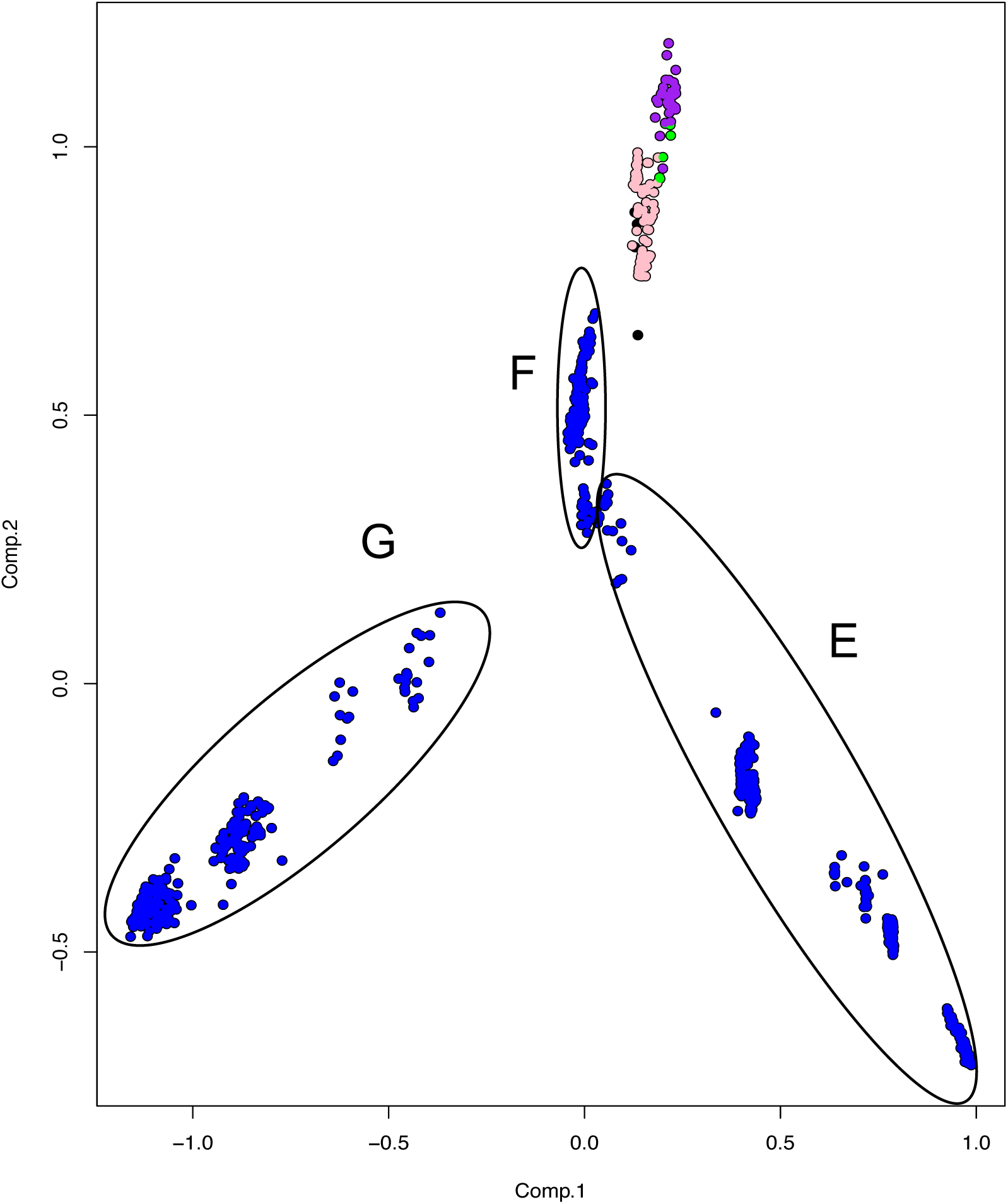
Principal component analysis (PCA) of *Mycobacterium bovis* lineages. Principal Component Analysis constructed using the SNP (single nucleotide polymorphism) matrix of the 823 *Mycobacterium bovis* genomes. The four inferred *M. bovis* lineages are shown in purple - Lb1; green - Lb2; pink - Lb3; blue - Lb4; uncharacterized lineage - black. Circles represent *M. bovis* genomes emerging from the described nodes (E, F, G) of the phylogenetic tree depicted in Figure 1.

### Evolutionary scenario for the origin of *M. bovis*

Recent analyses point towards the evolutionary rate defined by Bos et al. (Bos et al., 2014) as being the most plausible to explain the trajectory of MTBC (Menardo et al., 2019; O’Neill et al., 2019). Our posterior Bayesian estimate indicates that the time of the most recent common ancestor (MRCA) for extant *M. bovis* lineages is 430 - 2,394 years BP (Before Present) (Figure 4). This maximum time obtained (2,394 years BP) (Figure 4) agrees with the archeological finding of *M. bovis* (defined by the loss of RD4) causing infection in four human skeletons from the Iron Age in South Siberia, with carbon-dating placing them within the 1,761 - 2,199 years BP range (Taylor et al., 2007). Interestingly, these ancient *M. bovis* strains do not have the deletion marker of CC Eu1 (i.e. RD17), serving as supportive evidence that Lb4 is indeed a recently-evolved lineage of *M. bovis*. The oldest description of tubercle (‘phymata’) lesions in cattle (and also in sheep and pigs) was done by the Greek physician Hippocrates (2,480-2,390 BP) (Herzog, Basel, 2003), also overlapping our maximum estimate for the MRCA.

Considering the upper-end time of 2,394 years BP, the predicted origin of the extant *M. bovis* lineages overlaps with the Ancient Roman period (2645-1544 BP). The paucity of *M. bovis* genomes from Africa, particularly East, West and North of Africa, precludes us to rule out an African origin of *M. bovis*. However, as most extant Lb1 and Lb2 strains are from Europe and Africa, we speculate that the beginning of *M. bovis* genetic diversification may have had an important contribution from ancient Romans, possibly due to their influence on animal husbandry practices in the Mediterranean and North of Africa. We should not neglect the fact that the world saw a demographical explosion and a deep connection between different cultures and communities during the Ancient Roman period (Jongman, 2012; Hanson et al., 2017). It has been suggested that this demographic explosion and economic rise greatly contributed to the spread of human TB worldwide, with naïve human populations from rural areas being exposed to the disease when moving to larger cities (Herzog, Basel, 2003; O’Neill et al., 2019). The economical flourishing was followed by a steep rise in agriculture and livestock capacity (Jongman, 2012). Archeological findings related to changes in cattle body size between Neolithic times and the Roman period (Meadow, 1988, 1989; Meadow and Zeder, 2004; Trentacoste et al., 2018) suggest the occurrence of modifications in cattle herding management in the Roman period, including improved feeding practices, probably aimed at increasing production capacity for draft, meat and/or milk (Kron, 2002). Reasons for breeding cattle varied considerably throughout the Roman territory and beyond due to culture traditions, diet preferences and climate conditions (Itan et al., 2009; Valenzuela-Lamas and Albarella, 2017). Nevertheless, by intensifying production systems and animal trade, which was most likely not centered solely in Roman territory, bovine TB may have dispersed significantly.

We cannot assume for certain that the disease, bovine TB, exclusively originated during the Ancient Roman period. The disease may have been present in cattle before this time, although caused by a different ancestor of the extant *M. bovis* lineages. It is still unclear if the ancestor of MTBC was a specific human pathogen or a generalist microorganism able to infect multiple host species (Brites et al., 2018). In the latter case, the intensification of the livestock system and increase in animal and human population density may have selected for pathogens either more adapted to animals (*M. bovis*) or more adapted to human beings (*M. tuberculosis*) over evolutionary time. It should be noted that genomes from representative *M. bovis* isolates from Eastern and Western Asia have yet to be sequenced and analyzed. The lack of information from that continent makes it impossible to fully explain the finding of an ancient human infection near present Mongolian border in Russia (1,761 - 2,199 years BP) (Taylor et al., 2007). Alternatively, the well-known commercial trade between the Roman Empire and Asia may also account for this dispersal.

### Bovine TB after the 16^th^ century

Our dating estimates and evolutionary predictions reveal a complex relationship between spatial dispersal and expansion of *M. bovis*. Bovine TB worldwide distribution is certainly influenced by import and export of cattle breeds over time. Non-European countries imported specialized breed cattle from single sources and then exported to other countries, while others imported animals from multiple locations. Estimates show that the MRCAs of extant *M. bovis* isolates occurring in Northern Ireland date between 54 to 325 BP. Archeological data from cattle specimens found in the UK (United Kingdom) indicates that these animals had a shared ancestry with wild British aurochs, being unrelated to the modern European taurine (Orlando, 2015). Accordingly, in Northern Ireland, cattle have been the mainstay of farming since the neolithic period (about 6,000 BP). Therefore, despite the ancient presence of cattle in these regions, the introduction of the extant *M. bovis* lineage Lb4 seems to be a more recent event, possibly due to the importation of taurine cattle from other areas of Europe.

Most of the USA isolates clustered in Lb4 (186/233), a few in Lb3 (40/233) and only three in Lb1 (3/233). A similar trend is observed in Mexico, with the majority of the isolates clustered in Lb4 (175/192) and fewer representatives in Lb3 (17/192). Our dating estimates are consistent with bovine TB introduction into these countries during the New World colonization period, with Lb3 dating from 137 to 906 BP (USA) and from 41 to 241 BP (Mexico), and Lb4 dating from 71 to 411 BP (USA) and 216 to 1151 BP (Mexico). The first cattle to be introduced into the USA came from the Iberian Peninsula in the 16^th^ century. This introduction was followed by cattle importation from Mexico (which also received animals from the Iberian Peninsula) and from England (Martínez et al., 2012). This is consistent with the introduction of Lb3 (CC Eu2) and Lb4 (CC Eu1), common lineages in the Iberian Peninsula and the UK, respectively. Isolates from USA and Mexico frequently clustered together, consistent with the close cattle trade relationship between these two countries throughout modern history.

At least two main introductions of Lb4 may have happened in New Zealand. The first is estimated to have occurred between 92 and 488 BP, while the second from 60 to 333 BP. Breeds of cattle were introduced into New Zealand in the 19^th^ century (Livingstone et al., 2015), which is within our time period range and suggests consequent spreading to wildlife. In contrast, *M. bovis* genomes from South Africa appeared clustered in Lb3 and Lb4 (Smith et al., 2011). Despite the small sample size, they are not present in the most ancient lineages, Lb1 and Lb2. In fact, previous genotyping studies have shown that the most common *M. bovis* CC in South Africa is Eu1, i.e. Lb4. Thus, in agreement with previous hypotheses (Michel et al., 2009; Smith et al., 2011), it is likely that bovine TB was introduced into South Africa following European colonization, and subsequently spilled over to naïve wildlife, with destructive consequences (Michel et al., 2006), as exemplified by *M. bovis* genomes from kudus, lions and buffalos analyzed herein.

### Concluding remarks

We propose the existence of at least four global lineages of *M. bovis*, named Lb1 and Lb2, occurring mostly in Africa and Europe, Lb3 present mainly in the Americas, Europe and South Africa, and Lb4 dispersed worldwide. Because these lineages tend to cluster based on geographical location rather than host species, it reinforces the idea that bovine TB eradication will only be attained once the disease is controlled in wildlife and vice-versa. The observed lack of host specificity supports the hypothesis of *M. bovis* being a generalist member of MTBC. Nonetheless, *M. bovis* ability to be transmitted among cattle is the main reason why this pathogen has spread geographically over evolutionary time, because of animal trade.

CC and/or SNP markers are proposed for each lineage. Our results shown that CC markers are only stable for Lb2 (Af1) and Lb4 (Eu1), while Lb1 and Lb3 can be better identified using whole genome sequencing and a provisional set of SNPs. The lack of stable CC markers in the most ancient lineage, Lb1, indicates that these pathogens have been the least studied, and that there is an urgent need for additional evaluations of *M. bovis* isolates from Africa, as well as from Asia (given the low number of genome representatives from this continent). Further sequencing of *M. bovis* isolates throughout the world will provide the opportunity to refine the identification of SNP and deletion markers specific of each lineage, as well as provide accurate data from geographic areas not explored in this study.

Our results delineate independent evolutionary trajectories of bacterial subpopulations (i.e. lineages) of *M. bovis* underlying the current disease distribution. Whether or not these events are associated with further specialization of *M. bovis* to the bovine species and breeds or increased/decreased virulence of this pathogen in domesticated and/or wildlife have yet to be determined. Lineages of *M. tuberculosis* are known for their virulence variations, warranting further similar studies regarding *M. bovis* lineages.

Finally, our dating estimates for the MCRA of extant *M. bovis* lineages agreed with previous studies based on independent archeological data (South Siberian skeletons at ∼2,000 BP). The results suggest that dispersion and expansion of the species may have increased during the Ancient Roman period, stimulated by the demographic expansion and rising need for cattle breeding. This may have contributed to an increased number of TB cases in humans, albeit of zoonotic origin. Subsequently, the history of economic trade, especially involving animals, may explain the continuous spread of bovine TB worldwide. By understanding the evolutionary origin and genomic diversification of *M. bovis*, we expect that the results presented herein will help pave the way to avoid future outbreaks of the disease in cattle, wildlife and humans.

## Acknowledgements

The authors are in debt to Gisele Oliveira de Souza and Carolina Bertelli de Souza Ferreira from the University of São Paulo, São Paulo, Brazil for invaluable technical assistance. We also thank the LaCTAD, UNICAMP, Campinas, Brazil for aiding in the genome sequencing of *M. bovis* isolates, and CEFAP, University of São Paulo, for computer core services. We are in debt with Dr Paulo Eduardo Brandão for continuous mentoring support throughout this study.

## Author Contributions

CZ: Performed and/or participated in all experiments, analyzed and interpreted the data, and wrote the manuscript. JP: Designed and performed dating divergence experiments, analyzed and interpreted the resulting data. ACG: Performed experiments involving the generation and analyses of the SNP matrix. RS: Designed and supervised specific bioinformatics analyses. TP: Provided bioinformatic assistance for the phylogenomic analysis. NC: Provided bioinformatics assistance for the phylogenomic analysis. AF: Performed experiments involving bacterial isolation and DNA extraction. CI: Performed experiments involving bacterial isolation and DNA extraction. JN, JS, MH: Designed experiments and analyzed the data. AMG: Designed and coordinated the study, analyzed the data and wrote the manuscript.

## References

Berg, S., Garcia-Pelayo, M. C., Müller, B., Hailu, E., Asiimwe, B., Kremer, K., et al. (2011). African 2, a clonal complex of *Mycobacterium bovis* epidemiologically important in East Africa. J. Bacteriol. 193, 670–8. doi:10.1128/JB.00750-10.

Bolger, A. M., Lohse, M., and Usadel, B. (2014). Trimmomatic: a flexible trimmer for Illumina sequence data. Bioinformatics 30, 2114–2120. doi:10.1093/bioinformatics/btu170.

Bos, K. I., Harkins, K. M., Herbig, A., Coscolla, M., Weber, N., Comas, I., et al. (2014). Pre-Columbian mycobacterial genomes reveal seals as a source of New World human tuberculosis. Nature 514, 494–7. doi:10.1038/nature13591.

Brites, D., Loiseau, C., Menardo, F., Borrell, S., Boniotti, M. B., Warren, R., et al. (2018). A New Phylogenetic Framework for the Animal-Adapted *Mycobacterium tuberculosis* Complex. Front. Microbiol. 9, 2820. doi:10.3389/fmicb.2018.02820.

Bruning-Fann, C., Robbe-Austerman, S., Kaneene, J., Thomsen, B., Jr, J. T., Ray, J., et al. (2017). Use of whole-genome sequencing and evaluation of the apparent sensitivity and specificity of antemortem tuberculosis tests in the investigation of an unusual outbreak of Mycobacterium bovis infection in a Michigan dairy herd. 251, 206–216. doi:10.2460/javma.251.2.206.

Casali, N., Nikolayevskyy, V., Balabanova, Y., Harris, S. R., Ignatyeva, O., Kontsevaya, I., et al. (2014). Evolution and transmission of drug-resistant tuberculosis in a Russian population. Nat. Genet. 46, 279–86. doi:10.1038/ng.2878.

Cingolani, P., Platts, A., Wang, L. L., Coon, M., Nguyen, T., Wang, L., et al. (2012). A program for annotating and predicting the effects of single nucleotide polymorphisms, SnpEff. Fly (Austin). 6, 80–92. doi:10.4161/fly.19695.

Coscolla, M., and Gagneux, S. (2014). Consequences of genomic diversity in Mycobacterium tuberculosis. Semin. Immunol. 26, 431–444. doi:10.1016/j.smim.2014.09.012.

Davidson, J. A., Loutet, M. G., O’Connor, C., Kearns, C., Smith, R. M. M., Lalor, M. K., et al. (2017). Epidemiology of *Mycobacterium bovis* Disease in Humans in England, Wales, and Northern Ireland, 2002-2014. Emerg. Infect. Dis. 23, 377–386. doi:10.3201/eid2303.161408.

de Jong, B. C., Adetifa, I., Walther, B., Hill, P. C., Antonio, M., Ota, M., et al. (2010). Differences between tuberculosis cases infected with *Mycobacterium africanum*, West African type 2, relative to Euro-American Mycobacterium tuberculosis: an update. FEMS Immunol. Med. Microbiol. 58, 102–5. doi:10.1111/j.1574-695X.2009.00628.x.

de la Fuente, J., Díez-Delgado, I., Contreras, M., Vicente, J., Cabezas-Cruz, A., Tobes, R., et al. (2015). Comparative Genomics of Field Isolates of *Mycobacterium bovis* and *M. caprae* Provides Evidence for Possible Correlates with Bacterial Viability and Virulence. PLoS Negl. Trop. Dis. 9, 1–22. doi:10.1371/journal.pntd.0004232.

Dippenaar, A., Parsons, S. D. C., Miller, M. A., Hlokwe, T., Gey van Pittius, N. C., Adroub, S. A., et al. (2017). Progenitor strain introduction of *Mycobacterium bovis* at the wildlife-livestock interface can lead to clonal expansion of the disease in a single ecosystem. Infect. Genet. Evol. 51, 235–238. doi:10.1016/J.MEEGID.2017.04.012.

Eldholm, V., Monteserin, J., Rieux, A., Lopez, B., Sobkowiak, B., Ritacco, V., et al. (2015). Four decades of transmission of a multidrug-resistant *Mycobacterium tuberculosis* outbreak strain. Nat. Commun. 6, 7119. doi:10.1038/ncomms8119.

Firdessa, R., Berg, S., Hailu, E., Schelling, E., Gumi, B., Erenso, G., et al. (2013). Mycobacterial lineages causing pulmonary and extrapulmonary tuberculosis, Ethiopia. Emerg. Infect. Dis. 19, 460–3. doi:10.3201/eid1903.120256.

Ford, C. B., Lin, P. L., Chase, M. R., Shah, R. R., Iartchouk, O., Galagan, J., et al. (2011). Use of whole genome sequencing to estimate the mutation rate of *Mycobacterium tuberculosis* during latent infection. Nat. Genet. 43, 482–486. doi:10.1038/ng.811.

Galagan, J. E. (2014). Genomic insights into tuberculosis. Nat. Rev. Genet. 15, 307–320. doi:10.1038/nrg3664.

Garnier, T., Eiglmeier, K., Camus, J., Medina, N., Mansoor, H., Pryor, M., et al. (2003). The Complete Genome Sequencing of *Mycobacterium bovis*. Proc. Natl. Acad. Sci. 100, 7887–82. doi:10.1073/pnas.1130426100.

Ghebremariam, M. K., Hlokwe, T., Rutten, V. P. M. G., Allepuz, A., Cadmus, S., Muwonge, A., et al. (2018). Genetic profiling of *Mycobacterium bovis* strains from slaughtered cattle in Eritrea. PLoS Negl. Trop. Dis. 12, e0006406. doi:10.1371/journal.pntd.0006406.

Hanson, J. W., Ortman, S. G., and Lobo, J. (2017). Urbanism and the division of labour in the Roman Empire. J. R. Soc. Interface 14. doi:10.1098/rsif.2017.0367.

Herzog, Basel H. (2003). History of Tuberculosis. Respiration 65, 5–15. doi:10.1159/000029220.

Hoang, D. T., Vinh, L. S., Flouri, T., Stamatakis, A., von Haeseler, A., and Minh, B. Q. (2018). MPBoot: fast phylogenetic maximum parsimony tree inference and bootstrap approximation. BMC Evol. Biol. 18, 11. doi:10.1186/s12862-018-1131-3.

Itan, Y., Powell, A., Beaumont, M. A., Burger, J., and Thomas, M. G. (2009). The origins of lactase persistence in Europe. PLoS Comput. Biol. 5, 17–19. doi:10.1371/journal.pcbi.1000491.

Jongman, W. M. (2012). “Re-constructing the roman economy,” in The Cambridge History of Capitalism Volume I: The Rise of Capitalism: From Ancient Origins to 1848, eds. Larry Neal and Jeffrey G. Williamson (Cambridge: Cambridge University Press), 75–100. doi:10.1017/CHO9781139095099.004.

Kandler, J. L., Mercante, A. D., Dalton, T. L., Ezewudo, M. N., Cowan, L. S., Burns, S. P., et al. (2018). Validation of Novel *Mycobacterium tuberculosis* Isoniazid Resistance Mutations Not Detectable by Common Molecular Tests. Antimicrob. Agents Chemother. 62. doi:10.1128/AAC.00974-18.

Kay, G. L., Sergeant, M. J., Zhou, Z., Chan, J. Z.-M., Millard, A., Quick, J., et al. (2015). Eighteenth-century genomes show that mixed infections were common at time of peak tuberculosis in Europe. Nat. Commun. 6, 6717. doi:10.1038/ncomms7717.

Koboldt, D. C., Zhang, Q., Larson, D. E., Shen, D., McLellan, M. D., Lin, L., et al. (2012). VarScan 2: somatic mutation and copy number alteration discovery in cancer by exome sequencing. Genome Res. 22, 568–76. doi:10.1101/gr.129684.111.

Kohl, T. A., Utpatel, C., Niemann, S., and Moser, I. (2018). *Mycobacterium bovis* persistence in two different captive wild animal populations in Germany: A longitudinal molecular epidemiological study revealing pathogen transmission by whole-genome sequencing. J. Clin. Microbiol. 56, 1–9. doi:10.1128/JCM.00302-18d.

Kraemer Zimpel, C., Brandão, P. E., Souza Filho, A. F. de, De Souza, R. F., Ykuta, C. Y., Soares Ferreira Neto, J., et al. (2017). Complete genome sequencing of Mycobacterium bovis SP38 and comparative genomics of Mycobacterium bovis and M. tuberculosis strains. Front. Microbiol. 8, 2389. doi:10.3389/FMICB.2017.02389.

Kron, G. (2002). Archaeozoology and the Productivity of Roman Livestock Farming. Münstersche Beiträge zur Antiken Handel. 21, 53–73.

Lasserre, M., Fresia, P., Greif, G., Iraola, G., Castro-Ramos, M., Juambeltz, A., et al. (2018). Whole genome sequencing of the monomorphic pathogen *Mycobacterium bovis* reveals local differentiation of cattle clinical isolates. BMC Genomics 19, 1–14. doi:10.1186/s12864-017-4249-6.

Li, H. (2011). A statistical framework for SNP calling, mutation discovery, association mapping and population genetical parameter estimation from sequencing data. Bioinformatics 27, 2987–93. doi:10.1093/bioinformatics/btr509.

Li, H., and Durbin, R. (2010). Fast and accurate long-read alignment with Burrows–Wheeler transform. Bioinformatics 26, 589–595. doi:10.1093/bioinformatics/btp698.

Lillebaek, T., Norman, A., Rasmussen, E. M., Marvig, R. L., Folkvardsen, D. B., Andersen, Å. B., et al. (2016). Substantial molecular evolution and mutation rates in prolonged latent *Mycobacterium tuberculosis* infection in humans. Int. J. Med. Microbiol. 306, 580–585. doi:10.1016/J.IJMM.2016.05.017.

Livingstone, P. G., Hancox, N., Nugent, G., and de Lisle, G. W. (2015). Toward eradication: the effect of *Mycobacterium bovis* infection in wildlife on the evolution and future direction of bovine tuberculosis management in New Zealand. N. Z. Vet. J. 63 Suppl 1, 4–18. doi:10.1080/00480169.2014.971082.

Majoor, C. J., Magis-Escurra, C., van Ingen, J., Boeree, M. J., and van Soolingen, D. (2011). Epidemiology of *Mycobacterium bovis* Disease in Humans, the Netherlands, 1993–2007. Emerg. Infect. Dis. 17, 457–463. doi:10.3201/eid1703.101111.

Malone, K. M., and Gordon, S. V (2017). Strain Variation in the Mycobacterium tuberculosis Complex: Its Role in Biology, Epidemiology and Control. 1019. doi:10.1007/978-3-319-64371-7.

Mardia, K. V., Kent, J. T., and Bibby, J. M. (1979). Multivariate analysis. lLondon: Academic Press.

Martínez, A. M., Gama, L. T., Cañón, J., Ginja, C., Delgado, J. V., Dunner, S., et al. (2012). Genetic Footprints of Iberian Cattle in America 500 Years after the Arrival of Columbus. PLoS One 7, e49066. doi:10.1371/journal.pone.0049066.

Meadow, R. H. (1988). Archeology: The Archaeology of Animals. Simon J. M. Davis. Am. Anthropol. 90, 697. doi:10.1525/aa.1988.90.3.02a00220.

Meadow, R. H. (1989). “Osteological evidence for process of animal domestication,” in The Walking larder : patterns of domestication, pastoralism, and predation, 80–90.

Meadow, R., and Zeder, M. (2004). Approaches to Faunal Analysis in the Middle East.

Meikle, V., Bianco, M. V, Blanco, F. C., Gioffré, A., Garbaccio, S., Vagnoni, L., et al. (2011). Evaluation of pathogenesis caused in cattle and guinea pig by a *Mycobacterium bovis* strain isolated from wild boar. BMC Vet. Res. 7, 37. doi:10.1186/1746-6148-7-37.

Menardo, F., Duchene, S., Brites, D., and Gagneux, S. (2019). The molecular clock of *Mycobacterium tuberculosis*. bioRxiv https://doi.org/10.1101/532390. doi:cc.

Michel, A. L., Bengis, R. G., Keet, D. F., Hofmeyr, M., Klerk, L. M. de, Cross, P. C., et al. (2006). Wildlife tuberculosis in South African conservation areas: Implications and challenges. Vet. Microbiol. 112, 91–100. doi:10.1016/j.vetmic.2005.11.035.

Michel, A. L., Coetzee, M. L., Keet, D. F., Maré, L., Warren, R., Cooper, D., et al. (2009). Molecular epidemiology of Mycobacterium bovis isolates from free-ranging wildlife in South African game reserves. Vet. Microbiol. 133, 335–343. doi:https://doi.org/10.1016/j.vetmic.2008.07.023.

Müller, B., Hilty, M., Berg, S., Garcia-Pelayo, M. C., Dale, J., Boschiroli, M. L., et al. (2009). African 1, an epidemiologically important clonal complex of *Mycobacterium bovis* dominant in Mali, Nigeria, Cameroon, and Chad. J. Bacteriol. 191, 1951–60. doi:10.1128/JB.01590-08.

Nguyen, L.-T., Schmidt, H. A., von Haeseler, A., and Minh, B. Q. (2015). IQ-TREE: A Fast and Effective Stochastic Algorithm for Estimating Maximum-Likelihood Phylogenies. Mol. Biol. Evol. 32, 268–274. doi:10.1093/molbev/msu300.

O’Neill, M. B., Shockey, A., Zarley, A., Aylward, W., Eldholm, V., Kitchen, A., et al. (2019). Lineage specific histories of *Mycobacterium tuberculosis* dispersal in Africa and Eurasia. Mol. Ecol. 28, mec.15120. doi:10.1111/mec.15120.

Olea-Popelka, F., Muwonge, A., Perera, A., Dean, A. S., Mumford, E., Erlacher-Vindel, E., et al. (2017). Zoonotic tuberculosis in human beings caused by *Mycobacterium bovis* - a call for action. Lancet Infect. Dis. 17, e21–e25. doi:10.1016/S1473-3099(16)30139-6.

Orlando, L. (2015). The first aurochs genome reveals the breeding history of British and European cattle. Genome Biol. 16, 225. doi:10.1186/s13059-015-0793-z.

Orloski, K., Robbe-austerman, S., Stuber, T., Hench, B., and Schoenbaum, M. (2018a). Whole genome sequencing of *Mycobacterium bovis* isolated from livestock in the United States, 1989-2018. Front. Vet. Sci. 5:253, 1–23. doi:10.3389/fvets.2018.00253.

Orloski, K., Robbe-austerman, S., Stuber, T., Hench, B., and Schoenbaum, M. (2018b). Whole genome sequencing of Mycobacterium bovis isolated from livestock in the United States, 1989-2018. Front. Vet. Sci. 5:253, 1–23. doi:10.3389/fvets.2018.00253.

Patané, J. S. L., Martins, J., Castelão, A. B., Nishibe, C., Montera, L., Bigi, F., et al. (2017). Patterns and Processes of *Mycobacterium bovis* Evolution Revealed by Phylogenomic Analyses. Genome Biol. Evol. 9, 521–535. doi:10.1093/gbe/evx022.

Pepperell, C. S., Casto, A. M., Kitchen, A., Granka, J. M., Cornejo, O. E., Holmes, E. C., et al. (2013). The role of selection in shaping diversity of natural *M. tuberculosis* populations. PLoS Pathog. 9, e1003543. doi:10.1371/journal.ppat.1003543.

Portevin, D., Gagneux, S., Comas, I., and Young, D. (2011). Human Macrophage Responses to Clinical Isolates from the Mycobacterium tuberculosis Complex Discriminate between Ancient and Modern Lineages. PLoS Pathog. 7, e1001307. doi:10.1371/journal.ppat.1001307.

Price-Carter, M., Brauning, R., De-Lisle, G. W., Livingstone, P., Neill, M., Sinclair, J., et al. (2018). Whole Genome Sequencing for Determining the Source of *Mycobacterium bovis* Infections in Livestock Herds and Wildlife in New Zealand. Front. Vet. Sci. 5, 1–13. doi:https://doi.org/10.3389/fvets.2018.00272.

Razo, C. A. P., Hernández, E. R., Ponce, S. I. R., Suazo, F. M., Robbe-Austerman, S., Stuber, T., et al. (2018). Research article molecular epidemiology of cattle tuberculosis in mexico through whole-genome sequencing and spoligotyping. PLoS One 13, 1–14. doi:10.1371/journal.pone.0201981.

Robinson, D. F., and Foulds, L. R. (1981). Comparison of phylogenetic trees. Math. Biosci. 53, 131–147. doi:10.1016/0025-5564(81)90043-2.

Rodriguez-Campos, S., Schürch, A. C., Dale, J., Lohan, A. J., Cunha, M. V., Botelho, A., et al. (2012). European 2 – A clonal complex of *Mycobacterium bovis* dominant in the Iberian Peninsula. Infect. Genet. Evol. 12, 866–872. doi:10.1016/j.meegid.2011.09.004.

Sandoval-Azuara, S. E., Muñiz-Salazar, R., Perea-Jacobo, R., Robbe-Austerman, S., Perera-Ortiz, A., López-Valencia, G., et al. (2017). Whole genome sequencing of Mycobacterium bovis to obtain molecular fingerprints in human and cattle isolates from Baja California, Mexico. Int. J. Infect. Dis. 63, 48–56. doi:10.1016/j.ijid.2017.07.012.

Smith, N. H., Berg, S., Dale, J., Allen, A., Rodriguez, S., Romero, B., et al. (2011). European 1: A globally important clonal complex of *Mycobacterium bovis*. Infect. Genet. Evol. 11, 1340–1351. doi:10.1016/j.meegid.2011.04.027.

Suchard, M. A., Lemey, P., Baele, G., Ayres, D. L., Drummond, A. J., and Rambaut, A. (2018). Bayesian phylogenetic and phylodynamic data integration using BEAST 1.10. Virus Evol. 4. doi:10.1093/ve/vey016.

Taylor, G. M., Murphy, E., Hopkins, R., Rutland, P., and Chistov, Y. (2007). First report of *Mycobacterium bovis* DNA in human remains from the Iron Age. Microbiology 153, 1243–1249. doi:10.1099/mic.0.2006/002154-0.

Trentacoste, A., Nieto-Espinet, A., and Valenzuela-Lamas, S. (2018). Pre-Roman improvements to agricultural production: Evidence from livestock husbandry in late prehistoric Italy. doi:10.1371/journal.pone.0208109.

Valenzuela-Lamas, S., and Albarella, U. (2017). Animal Husbandry across the Western Roman Empire: Changes and Continuities. Eur. J. Archaeol. 20, 402–415. doi:10.1017/eaa.2017.22.

Vargas-Romero, F., Mendoza-Hernández, G., Suárez-Güemes, F., Hernández-Pando, R., and Castañón-Arreola, M. (2016). Secretome profiling of highly virulent *Mycobacterium bovis* 04-303 strain reveals higher abundance of virulence-associated proteins. Microb. Pathog. 100, 305–311. doi:10.1016/J.MICPATH.2016.10.014.

Venables, W. N., and Ripley, B. D. (2002). Modern Applied Statistics with S. New York, NY: Springer-Verlag New York doi:10.1007/978-0-387-21706-2.

Walker, T. M., Kohl, T. A., Omar, S. V, Hedge, J., Del Ojo Elias, C., Bradley, P., et al. (2015). Whole-genome sequencing for prediction of *Mycobacterium tuberculosis* drug susceptibility and resistance: a retrospective cohort study. Lancet. Infect. Dis. 15, 1193–1202. doi:10.1016/S1473-3099(15)00062-6.

Warren, R. M., Streicher, E. M., Sampson, S. L., van der Spuy, G. D., Richardson, M., Nguyen, D., et al. (2002). Microevolution of the Direct Repeat Region of Mycobacterium tuberculosis: Implications for Interpretation of Spoligotyping Data. J. Clin. Microbiol. 40, 4457–4465. doi:10.1128/JCM.40.12.4457-4465.2002.

Waters, W. R., Palmer, M. V, Thacker, T. C., Bannantine, J. P., Vordermeier, H. M., Hewinson, R. G., et al. (2006). Early antibody responses to experimental *Mycobacterium bovis* infection of cattle. Clin. Vaccine Immunol. 13, 648–54. doi:10.1128/CVI.00061-06.

Wedlock, D. N., Aldwell, F. E., Collins, D. M., de Lisle, G. W., Wilson, T., and Buddle, B. M. (1999). Immune responses induced in cattle by virulent and attenuated *Mycobacterium bovis* strains: correlation of delayed-type hypersensitivity with ability of strains to grow in macrophages. Infect. Immun. 67, 2172–7.

World Health Organization (2018). WHO Tuberculosis Report 2018.

Wright, D. M., Allen, A. R., Mallon, T. R., McDowell, S. W. J., Bishop, S. C., Glass, E. J., et al. (2013). Field-Isolated Genotypes of *Mycobacterium bovis* Vary in Virulence and Influence Case Pathology but Do Not Affect Outbreak Size. PLoS One 8, e74503. doi:10.1371/journal.pone.0074503.

Xia, E., Teo, Y.-Y., and Ong, R. T.-H. (2016). SpoTyping: fast and accurate in silico *Mycobacterium* spoligotyping from sequence reads. Genome Med. 8, 19. doi:10.1186/s13073-016-0270-7.

Zimpel, C. K., Brandão, P. E., de Souza Filho, A. F., de Souza, R. F., Ikuta, C. Y., Ferreira Neto, J. S., et al. (2017a). Complete Genome Sequencing of *Mycobacterium bovis* SP38 and Comparative Genomics of *Mycobacterium bovis* and *M. tuberculosis* Strains. Front. Microbiol. 8, 2389. doi:10.3389/fmicb.2017.02389.

Zimpel, C. K., Brum, J. S., de Souza Filho, A. F., Biondo, A. W., Perotta, J. H., Dib, C. C., et al. (2017b). *Mycobacterium bovis* in a European bison (*Bison bonasus*) raises concerns about tuberculosis in Brazilian captive wildlife populations: a case report. BMC Res. Notes 10, 91. doi:10.1186/s13104-017-2413-3.

